# Insertional oncogenesis by HPV70 revealed by multiple genomic analyses in a clinically HPV-negative cervical cancer

**DOI:** 10.1101/634857

**Authors:** Anne Van Arsdale, Nicole E. Patterson, Elaine C. Maggi, Lorenzo Agoni, Koenraad Van Doorslaer, Bryan Harmon, Nicole Nevadunsky, Dennis Y.S. Kuo, Mark H Einstein, Jack Lenz, Cristina Montagna

**Affiliations:** Departments of Obstetrics & Gynecology and Women’s Health, Albert Einstein College of Medicine, Yeshiva University, Bronx, NY, USA; Genetics, Albert Einstein College of Medicine, Yeshiva University, Bronx, NY, USA; Department of Women’s and Children’s Health, Obstetrics & Gynecology Unit, Fondazione Poliambulanza Istituto Ospedaliero, Brescia, Italy; School of Animal and Comparative Biomedical Sciences College of Agriculture and Life Sciences BIO5 Institute University of Arizona, AZ, USA; Pathology, Albert Einstein College of Medicine, Yeshiva University, Bronx, NY, USA; Department of Obstetrics, Gynecology, and Women’s Health, Rutgers New Jersey Medical School. Newark, NJ, USA

**Keywords:** HPV70, HPV DNA, hybridization capture, long range sequencing, fluorescent *in situ* hybridization, *BCL11B*, oncogene, insertional oncogenesis, cervical carcinoma, cancer.

## Abstract

Cervical carcinogenesis, the second leading cause of cancer death in women worldwide, is caused by multiple types of human papillomaviruses (HPVs). To investigate a possible role for HPV in a cervical carcinoma that was HPV-negative by PCR testing, we performed HPV DNA hybridization capture plus massively parallel sequencing. This detected a subgenomic, URR- E6-E7-E1 segment of HPV70 DNA, a type not generally associated with cervical cancer, inserted in an intron of the B-cell lymphoma/leukemia 11B (*BCL11B*) gene in the human genome. Long range DNA sequencing confirmed the virus and flanking *BCL11B* DNA structures including both insertion junctions. Global transcriptomic analysis detected multiple, alternatively spliced, HPV70-*BCL11B*, fusion transcripts with fused open reading frames. The insertion and fusion transcripts were present in an intraepithelial precursor phase of tumorigenesis. These results suggest oncogenicity of HPV70, identify novel *BCL11B* variants with potential oncogenic implications, and underscore the advantages of thorough genomic analyses to elucidate insights into HPV-associated tumorigenesis.

**Statement of Significance:** Multiple HPV types have been defined as high risk for cancer causation. However, genomic analyses applied here detected a non-high risk HPV in a carcinoma that was HPV negative, and elucidated virally-associated tumorigenic genetic events. This stresses the importance of thorough genomic analyses for elucidating genetic processes in HPV-associated tumorigenesis.

**Author Summary:** Cervical cancer is the second leading cause of cancer death in women worldwide. Most cervical cancers are caused by one of 15 high risk types of human papilloma viruses (HPVs), although hundreds of types of HPVs exist. We used a series of contemporary genomics analyses to examine a cervical cancer that was clinically determined to be HPV-negative. These detected DNA of HPV70, an HPV type not considered to be high risk, in the tumor. Approximately half of the HPV70 DNA genome was present including the viral E6 and E7 oncogenes. Moreover, the viral DNA was inserted into the *BCL11B* gene in the human genome. *BCL11B* is known to be mutated in certain human cancers. The HPV70 DNA interacted with the human *BCL11B* gene to produce altered forms of RNA encoding unusual, truncated forms of the *BCL11B* protein. These results strongly implicate HPV70 as being oncogenic, suggest that this tumor was caused by a combination of viral oncogenes plus the virally-activated human *BCL11B* gene, demonstrate novel truncated *BCL11B* variants with oncogenic implications, and underscore the advantages of thorough genomic analyses to elucidate HPV tumorigenesis insights

## Introduction

Infections with HPV types that are classified as high-risk by epidemiological criteria (hrHPVs) underlie the vast majority of invasive cervical carcinomas (ICCs) (1). ICCs are responsible for 4.5% of cancer deaths in women worldwide and comprise a substantial fraction of the estimated 12% of cancers worldwide caused by various viruses (2). Multiple large studies showed hrHPV types to be present in at least 90% of ICC’s using established, PCR- or hybridization-based methods for viral DNA detection (3, 4). HPV types categorized as Group 1 known carcinogens by the International Agency for Research on Cancer (IARC) working group are phylogenetically classified within the high-risk *Alphapapillomavirus*-7 and *Alphapapillomavirus*-9 clades of the *Alphapapillomavirus* genus and include the HPV types HPV16, HPV18, HPV31, HPV33, HPV35, HPV39, HPV45, HPV51, HPV52, HPV56, HPV58, and HPV59, with HPV16 and HPV18 being the most frequently detected types in ICC (5). Despite also belonging to the species *Alphapapillomavirus* 7 and *Alphapapillomavirus* 9, other less frequent HPV types also belonging to these clades (HPV26, HPV30, HPV34, HPV53, HPV66, HPV67, HPV69, HPV70, HPV73,HPV82 and HPV85), do not fulfill the rigorous epidemiological criteria required to be categorized as Group 1 carcinogens (5). At least 40 HPV types are known to infect the urogenital tract (1, 6, 7). Up to ~10% of invasive cervical cancers are currently considered HPV-negative (3, 4). This might be because of tissue sampling errors, the portion of the HPV genome targeted by the assay was deleted, or the tumor is associated with a low prevalence HPV type (8). An additional formal possibility is that some small fraction has a non-HPV etiology.

Population screening for women for cervical cancer historically has been accomplished by cytology (Papanicolaou test), but multiple clinical trials have provided evidence of improved sensitivity of hrHPV testing for detection of cervical cancer precursor lesions compared to cytology alone over multiple rounds of screening and follow up (9–13). The HPV types included in the screening assays are based on epidemiological prevalence data that balance sensitivity and specificity so as not to over-triage women to secondary screening such as colposcopy, and they currently include at most the 14 hrHPV types. Exclusion of less prevalent types from HPV screening reduces possible iatrogenic morbidity from subsequent colposcopy and excisional procedures following detection of viral types of uncertain pathological significance, and also reduces the costs involved in more extensive testing (14). Unlike population based screening, diagnostic HPV testing of identified pre-invasive and invasive cervical lesions is applicable to research driven assays.

While at least half of sexually active women incur a genital tract HPV infection during their lifetime (15), cervical tumorigenesis occurs in only a very small fraction of infected women. Tumorigenesis proceeds in a small fraction of women through a series of histopathologically defined, cervical intraepithelial neoplasia (CIN) lesions of progressively decreasing likelihood, CIN1, CIN2, and CIN3. The vast majority of HPV infections resolve spontaneously, with even most CIN lesions resolving spontaneously and only 20% to 30% of CIN3 lesions progressing to life-threatening ICC.

HPV infection proceeds with the ~8 kbp viral DNA genome replicating as a circular episome initially in the basal epithelial cell layer. A key step in HPV-associated tumorigenesis is integration of viral DNA into the human genome (16–19). At least part of the viral genome is inserted into human DNA in most HPV-induced ICC’s, and the integration process is thought to be mediated by host DNA repair mechanisms. Integration causes stable association of the virally-encoded oncogenes with a host cell, triggers human genome rearrangements, and drives expression of the human oncogenes that flank the sites of integration (20–25). In most ICC’s, the HPV genome is inserted near one allele of a human, dominant-acting oncogene, and many human oncogenes have been identified as recurring HPV insertion sites in tumors (22, 26–30). The presence of viral DNA adjacent to such human oncogenes is thought to be a consequence of integration occurring at a stochastic position within the human genome, followed by selectively advantageous, clonal proliferation of cells that suffered an integration near such oncogenes. Insertional oncogenesis by activation of flanking oncogenes is a long studied and thoroughly established mechanism of tumorigenesis by various retroviruses in experimental and naturally occurring animal systems in which insertion of viral DNA containing strong transcriptional elements, particularly enhancers, alters the transcription of the flanking gene (31–35). This is also the causal mechanism in the tumors that occurred during gene therapy studies in humans (36, 37). HPV activation of nearby oncogene transcription may entail a variety of mechanisms including fusion transcripts. These initiate at viral promoters and proceed into the host genes. Viral enhancers may also activate flanking oncogene promoters (38–40).

Approaches to detect integrated viral DNA have included RNA sequencing (RNAseq) to detect fusion transcripts, and DNA sequencing strategies including PCR or hybridization capture to identify the junctions between inserted HPV and human genome DNA (20–23). However, in most instances only one of the two junctions between the linear HPV DNA fragment and human DNA was identified, and structural characterization of the inserted viral DNA was incomplete or inferred. Here we describe the use of a multiple HPV type, hybridization capture approach yielding sufficient DNA recovery and massively parallel sequencing depth to identify both junctions of an inserted HPV DNA. Combined with additional molecular genetic techniques, this allows comprehensive, robust characterization of HPV cervical tumorigenesis.

## Results

HPV DNA analysis was applied to a cervical carcinoma chosen specifically based on its HPV-negative, clinical testing status at the time of cancer diagnosis. A 46-year-old woman presented at the emergency room of our institution with International Federation of Gynecology and Obstetrics (FIGO) stage IIB, squamous cell ICC. HPV testing using the *cobas* test (Roche Molecular Systems), which detects 14 hrHPV types by PCR of a viral L1 gene segment followed by probe hybridization, was negative. Tumor DNA was subjected to hybridization capture enrichment of HPV DNA followed by massively parallel, next generation, DNA sequencing (HC+NGS) to search for any HPV DNA present in the lesion and, if so, determine whether it was inserted into the human genome. Targeted enrichment was accomplished using hybridization to a custom designed capture probe set containing DNA from 15 different hrHPV types (HPV 6, HPV11, HPV16, HPV18, HPV31, HPV33, HPV35, HPV39, HPV45, HPV52, HPV56, HPV58, HPV59, HPV68, HPV69). Each probe consists of a biotinylated, roughly 150 nucleotide long DNA strand specific for each HPV type (about 55 probes per HPV genome with probe overlap) encompassing the positive strands of the full ~8 kbp HPV genomes to allow capture along each entire viral genome.

HC+NGS of the tumor DNA yielded 155,832 unique paired end reads that mapped to the HPV70 genome (Figure 1), with average coverage depth of 1054X. All reads mapped to one 3980 bp segment encompassing about half of the viral genome from positions 6306-2380 including the entire upstream regulatory region (URR), E6 and E7 segments plus parts of the E1 and L1 genes. The segment contained 55 single nucleotide substitutions (1.4%) relative to the HPV70 reference sequence (**Supplementary Figure 1**). While HPV70 was not specifically targeted by the probes used, the HPV70 DNA fragments were presumably recovered by hybridization to related species *Alphapapillomavirus 7* types in the set, most likely HPV39 and HPV68. Forced alignment to a secondary custom genome containing only the HPV types included in the hybridization probe set was attempted, however alignment in this setting was both lower in unique reads (72,681 aligned read pairs to HPV type 68) and identity(<90%). More importantly, detection of HPV70 DNA demonstrated that this clinically HPV-negative tumor in fact did contain HPV, albeit from an uncommon type currently not classified as high risk.

**Figure 1.**
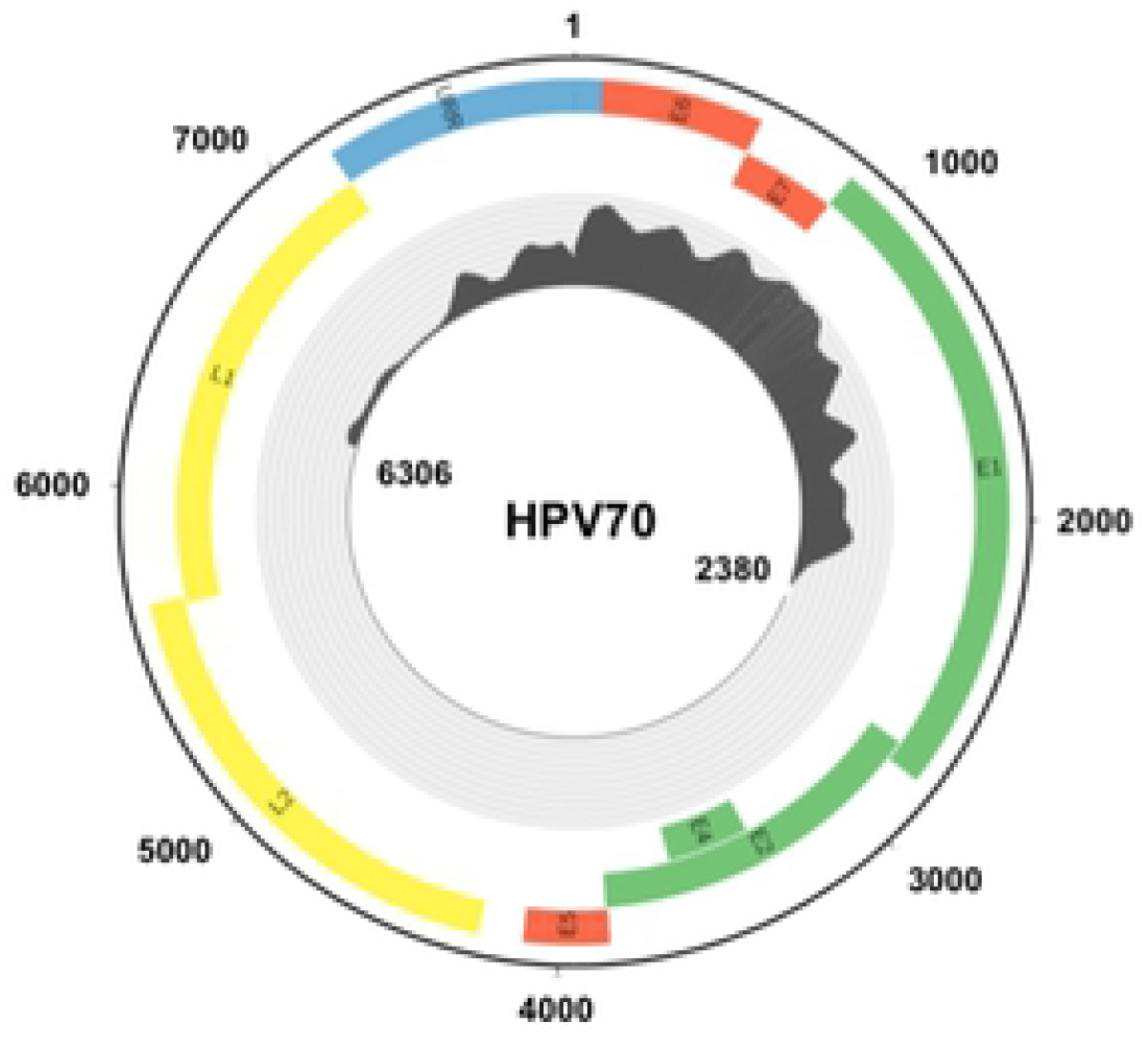
HPV70 genome coverage of sequence reads from viral DNA hybridization capture plus Illumina short read sequencing. The outer circle shows the circular episomal form of the HPV70 genome. Defined HPV oncogenes are in red, other early genes in green, late genes in yellow, and the upstream regulatory region (URR) in blue. The inner circle shows a histogram of sequencing coverage ranging from of 0 to 4200 reads. All reads corresponded to the segment from genome positions 6906 to 2380 containing the URR E6 and E7, with no reads detected in the remaining half of the viral genome.

The HC+NGS analysis detected junctions between the ends of the viral segment and sequences in the last intron of the human *BCL11B* gene between the third and fourth exons (Figure 2A). *BCL11B* encodes a Kruppel-like zinc finger protein that was previously identified as an integration site of HPV16 in at least two ICC’s (41, 42) and is mutated in other human cancers, most notably T-lymphocyte tumors (43–47). The occurrence of the two junctions at nearby genomic positions in *BCL11B* suggested that the 3980 bp HPV70 segment comprised a single insertion at this site (Figure 2A). To confirm the implied structure of HPV70 in a *BCL11B* allele, the tumor was subjected to intensive molecular genetic characterization by long range DNA sequencing, short range whole genome sequencing (WGS), Affymetrix Genome-Wide Human SNP Array 6.0 analysis, and RNA sequencing (RNAseq).

**Figure 2.**
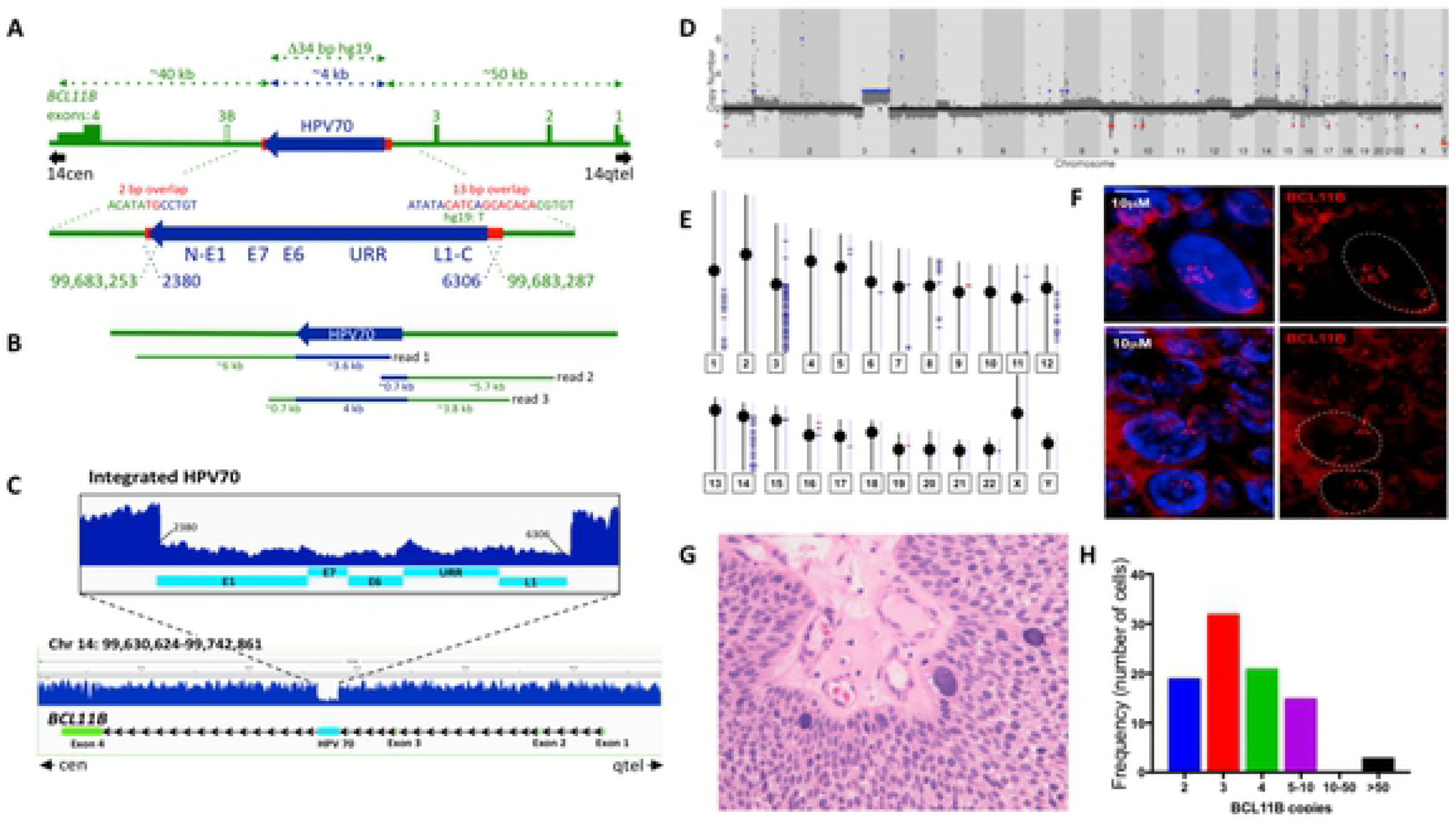
DNA sequencing analysis of the HPV70 DNA-containing tumor. **A)** Genetic diagram of the inferred structure of the HPV70 DNA segment insertion into an allele of the *BCL11B* gene based on hybridization capture and Illumina sequencing of HPV70 DNA. HPV70 components and genome positions are in blue. *BCL11B* components and positions on chromosome 14 (hg19) are in green. Red shows the positions and sequences of micro-homology at the junctions between human DNA and the viral DNA insert. Approximate distances of components are depicted by the dotted arrows at the top. Green rectangles and numbers indicate *BCL11B* exons with the 5’ and 3’ untranslated regions at lower height. Exons and HPV70 DNA are drawn larger than the scale of the full-length *BCL11B* region. **B)** Genome coverage and approximate sizes of the Oxford Nanopore MinION sequence reads encompassing HPV70 DNA confirming the single ~4 kbp insert and the structure of the immediately surrounding *BCL11B* sequences. **C)** Illumina short read, whole genome sequence coverage of the *BCL11B* gene on chromosome 14. The full-length *BCL11B* gene is at the bottom. An enlarged view of the HPV70 segment and immediately surrounding human DNA showing the lower coverage of the HPV70 segment is presented above. **D)** Gingkos plot of whole genome sequencing depth across the human genome from chromosome 1 through chromosome X. Red dots indicate chromosomal loss and blue dots indicates chromosomal gain. **E)** Human genome-wide Affymetrix SNP array coverage on each human chromosome. Short horizontal blue lines indicate segments where copy number ≥ 3 was detected. **F)** Fluorescence in situ hybridization of a BAC probe encompassing the *BCL11B* gene to a 10 mM histological section from the tumor showing the *BCL11B* gene copy number variation among the tumor cells. Red is the fluorescently labeled BAC DNA probe comprising the *BCL11B* region. The top and bottom panels show two different fields. DAPI staining of nuclei in each is shown in the left panels and was filtered from the right panels. **G)** Hematoxylin and eosin stained histological section from the tumor showing several large cells and their variable sizes. **H)** Quantification of the number of *BCL11B* loci in individual tumor cells. The number of fluorescent signals in each of 90 cells as in Panel F was counted.

Long range DNA sequencing utilizing an Oxford Nanopore MinION flowcell yielded 3.56X haploid genome coverage including three reads encompassing the HPV70 DNA insertion in *BCL11B* (Figure 2B). This unambiguously confirmed the structure of the insertion. WGS at 60X average genome coverage also confirmed both virus-human junctions, the sequence of the HPV70 segment detected by HC+NGS, and the absence of the remainder of HPV70 (Figure 2C). The number of WGS read counts across HPV70 was about one-third the number across *BCL11B*, indicating that HPV70 DNA was present in one allele of the *BCL11B* gene, and that one or more normal *BCL11B* alleles were also present, which is consistent with the insertion allele acting dominantly in tumorigenesis. Human DNA copy number was assessed genome-wide by generating a Ginko plot of WGS reads across the human genome (Figure 2D). Gain of the entire chromosome 3q arm was detected, along with copy number increases on human chromosomes 1q, 5p, 8, 12q and 14q, the latter including the *BCL11B* locus. 3q trisomy, which is highly recurrent in ICC, has been proposed to define transition from cervical dysplasia to invasive carcinoma through effects of the telomerase reverse transcriptase encoding *TERT* gene (48, 49). The 1q, 5p, 8, 12q and 14q copy number variations were also detected by genome-wide SNP analysis (Figure 2E), thus confirming their presence in the tumor. The observation that they were not as pronounced as the 3q trisomy indicated that they were less uniform among the tumor cells and likely involved multiple segments of the cognate chromosomes.

To investigate the extent of variability of the *BCL11B* copy number among the tumor cells, fluorescent *in situ* hybridization (FISH) was performed on histological sections of the tumor using a BAC probe encompassing the *BCL11B* gene (Figure 2F). Quantification showed that most cells contained between two and ten *BCL11B* loci, with modal number three (Figure 2H). About 4% of cells had 50 or more copies (Figures 2F, 2H). These cells were very large compared to the other tumor cells (Figure 2F) and were evident in standard, hematoxylin plus eosin stained, histopathology sections (Figure 2G). In summary, there was extensive variability in *BCL11B* copy number among the tumor cells.

The only HPV70 DNA detected in the tumor was the 3980 bp insert in *BCL11B*. No full-length, circular viral genomes were detectable. The two viral-human DNA junctions encompassed short stretches of sequence identity or microhomology, 12/13 bp at one end and 2 bp at the other (Figure 2A). Insertion of HPV70 DNA was accompanied by deletion of 34 bp relative to the reference human genome. The viral DNA insertion allele comprised a minority fraction of all the *BCL11B* loci present in the tumor cell population (Figure 2C), with the *BCL11B* copy number being highly non-uniform among the tumor cells (Figure 2H).

Intronic insertion of the HPV70 viral segment including the URR and full length E6 and E7 genes in the same transcriptional orientation as *BCL11B* raised questions of whether both viral oncogenes and *BCL11B* were transcribed, and whether HPV70-*BCL11B* fusion transcripts were present. Therefore, RNAseq was performed on tumor RNA, and reads were aligned with the HPV70 segment-containing *BCL11B* gene (Figure 3A). HPV70 transcripts derived almost entirely from a segment encompassing E6, E7, and E1 precisely up to the standard E1^E4 5’ splice site (5’ss) (Figure 3B). E6 and E7 are normally translated from transcripts containing both open reading frames (ORFs) (50), and about 40% of the transcripts in this tumor were consistent with excision of the standard HPV E6* intron (Figure 3B). However, only few of the transcripts included the AUG start codon for E6 (Figure 3C), indicating that only a fraction of transcripts could contain the full-length E6 or E6* ORFs, and those that did had very short 5’ untranslated regions, implying that transcription initiated downstream of the standard early transcription start site used during early phase of the HPV replicative cycle, and suggesting that little if any E6 or E6* translation may have occurred in this tumor. In summary, the HPV RNAseq reads were consistent with most transcription initiating slightly downstream of where viral early transcription normally initiates during an HPV infection. Outside of the E6, E7, and E1 segment up to the E1^E4 5’ss, almost no HPV70 transcripts were detected (Figure 3B). E7, which is normally expressed from mRNAs containing both E6 and E7 ORFs, was the only viral ORF entirely covered by abundant transcripts. Moreover, the E6* splices that were detected must have occurred mostly in what is the 5’ UTR of E7 ORF-containing mRNAs in this tumor. During infections, HPVs encode the E4 ORF from transcripts with the E6 and E7 ORFs upstream of the E1^E4 5’ss that normally fuses the 5’ end of the E1 ORF to that of E4. The abrupt absence of viral transcripts downstream of the standard E1^E4 5’ss strongly suggested efficient splicing at this junction, but to a non-viral downstream 3’ss in this tumor.

**Figure 3.**
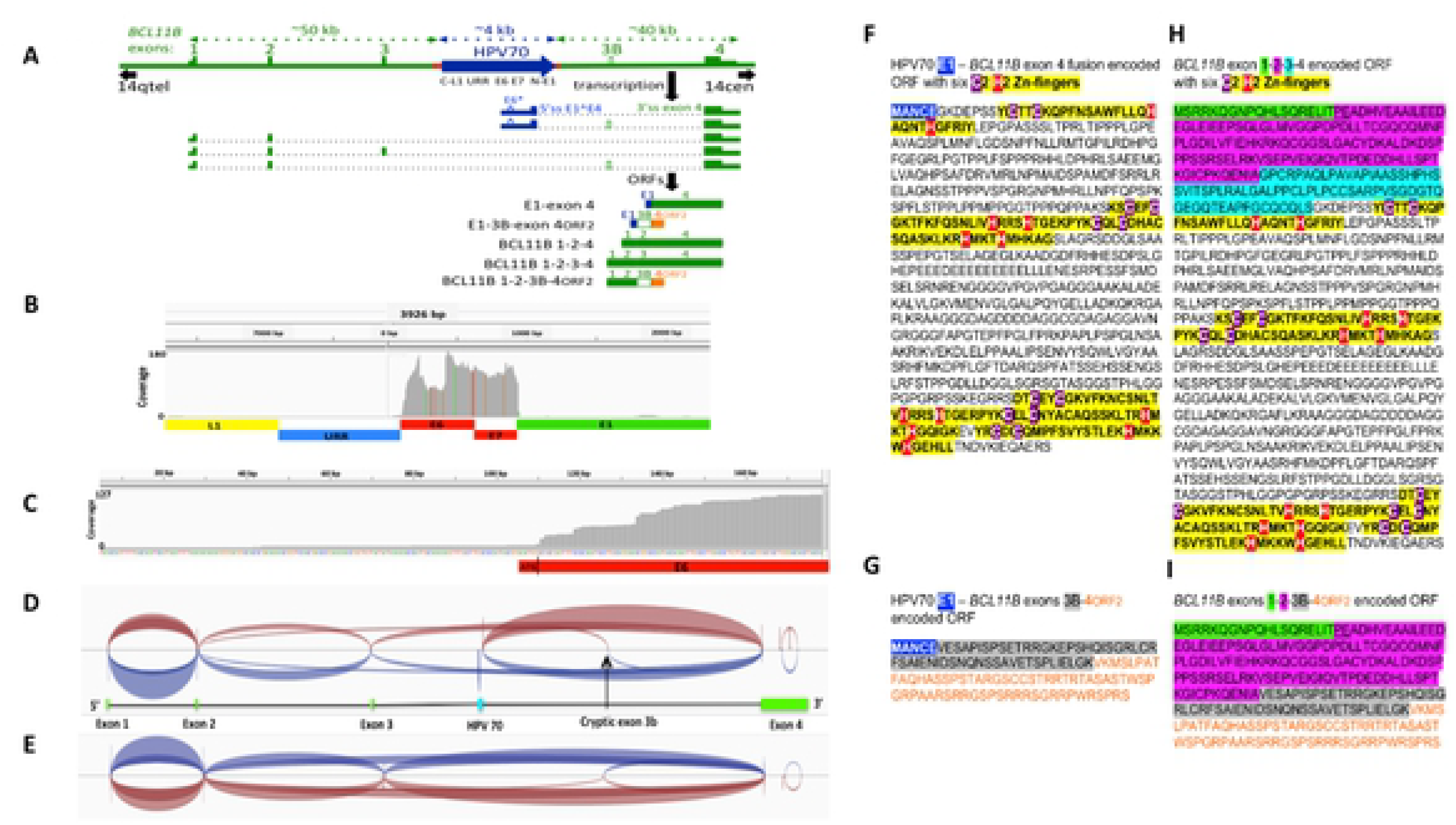
RNAseq analysis of HPV70 and *BCL11B* transcripts in the tumor. **A)** Genetic diagram of the *BCL11B* gene allele and the HPV70 DNA insert, with exons and HPV70 are drawn larger than scale of the full-length *BCL11B* gene segment. The 5’ to 3’ transcriptional orientation of both *BCL11B* and HPV70 is from left to right. Unspliced primary transcripts inferred from the 150 nucleotide RNAseq reads are shown below the genomic DNA with introns depicted as dotted segments. Positions of key splicing components are labeled, specifically the E1^E4 5’ss, the *BCL11B* exon 4 3’ss, and the E6* intron. Open reading frames (ORFs) encoded by the spliced mRNAs that were supported by split reads are diagramed below the primary transcripts. The top two are HPV70-*BCL11B* fusions, while the bottom three are *BCL11B* transcripts. HPV70 sequences are in blue, and *BCL11B* sequences are in green with variably spliced exon 3B in green outline and white center. The alternative ORF of exon 4 that is read from exon 3B-containing transcripts (ORF 2) is shown in orange. **B)** RNAseq coverage of the HPV70 segment showing that virtually all reads located to the region encompassing E6, E7 and the portion of E1 up to the E1^E4 5’ss. A decreased number of reads was present over the E6* intron indicating removal of that intron in a fraction of the transcripts. Colored vertical bars are nucleotides that differ from the HPV70 reference sequence. **C)** Enlarged view of RNAseq reads near the start codon for HPV70 E6 gene ORF showing the limited number of reads near the ATG start codon. The E6 ORF from the ATG start codon is shown in red at the bottom. **D)** Sashimi plot showing split read support for splice junctions. The diagram below the plot shows the positions of HPV70 DNA (blue) and the *BCL11B* exons (green), and represents the custom-generated genomic segment containing the HPV70 DNA insertion that was used to obtain the plot. **E)** Sashimi plot of *BCL11B* exon splice junctions obtained using the *BCL11B* genomic segment without the HPV70 DNA insertion demonstrating the inclusion of alternative transcripts with exon 3B in this tumor, including split reads supporting splicing from exon 2 to exon 3B. **F)** The amino acid sequence encoded by the HPV70 E1 – *BCL11B* exon 4 fusion ORF. HPV70 E1 encoded amino acids are highlighted in blue. The six C2H2 zinc fingers within the exon 4 encoded segment are highlighted in yellow with the cysteine residues in purple and the histidines in red. 66 split reads supported the E1 to exon 4 splice junction, five of which are presented in Supplementary Figure 2. **G)** The amino acid sequence encoded by the HPV70 E1 to *BCL11B* exon 3B and exon 4 fusion ORF. HPV70 E1 encoded amino acids are highlighted in blue. Exon 3B encoded amino acids are highlighted in gray. Exon 4 ORF2 encoded amino acids are shown in orange. The nucleotide sequence of exon 3B including intronic splicing signals along with the four split reads that support E1 to exon 3B splicing are shown in Supplementary Figure 3. **H)** Amino acid sequence of *BCL11B* encoded by exons 1, 2, 3 and 4 is shown for comparison. Exon 1 sequences are highlighted in green. Exon 2 sequences are highlighted in purple. Exon 3 sequences are highlighted in blue. the six zinc fingers encoded in exon 4 are highlighted as in Panel F. **I)** Inferred amino acid sequence encoded by *BCL11B* mRNA with alternative exon 3B. Segment colors match panels G and H. The two underlined amino acids (proline and glutamate) at the beginning of exon 2 in panels H and I show amino acids that are excluded by a known alternative splice junction between exons 1 and 2 of *BCL11B*.

Such splice junctions from HPV70 to *BCL11B* (Figure 3D) were identified in RNAseq split reads (**Supplementary Figure 2**). Most of these joined the viral E1^E4 5’ss to the 3’ss for *BCL11B* exon 4, showing that viral-host fusion transcripts were synthesized in this tumor. In addition, about 4% of E1^E4 5’ss’s were joined to a cryptic *BCL11B* exon, here termed exon 3B, between the viral insertion and exon 4 (Figure 3D). Cryptic exon 3B was spliced to exon 4 downstream, and both the 5’ and 3’ intronic sequences immediately flanking the cryptic exon had standard splice signals (**Supplementary Figure 3**). Alignment of the RNAseq reads with a normal *BCL11B* allele sequence without HPV70 identified all the standard splice junctions of the host gene, including variable inclusion of exon 3 (Figure 3E), indicating expression of normal *BCL11B* transcripts in the tumor as well, possibly from normal *BCL11B* alleles. In addition, splicing of a variant *BCL11B* transcript was identified in which the cryptic exon 3B was present between exons 2 and 4 (Figure 3E).

Transcripts with the most frequently detected splice junction (E1^E4 5’ss to 3’ss exon 4) encoded a fusion protein with the first five amino acids of HPV70 E1 fused to the portion of *BCL11B* encoding all six of its zinc fingers (Figure 3F). The E1^E4 splice to cryptic exon 3B joined the first five HPV70 E1 amino acids in frame to an ORF that completely spanned exon 3B, which in turn was spliced to exon 4 (Figure 3G). However, the splice junction from exon 3B to exon 4 caused exon 4 to be read in a different ORF than normal and encompassing 66 amino acids and no zinc fingers (Figure 3G). Similarly, while transcripts encompassing the standard *BCL11B* exons 1, 2, 3, and 4 encoded the standard six Zn-finger protein (Figure 3H), splicing of *BCL11B* transcripts from exon 2 to exon 3B fused the ORFs in frame, but the junction from exon 3B to exon 4 caused the same frameshift as in the viral fusion transcripts (Figure 3I). In summary, the HPV70 DNA insertion resulted mainly in HPV70-*BCL11B* fusion transcripts encoding an unusual, N-terminally truncated form of the *BCL11B* oncogene with five HPV70 E1 amino acids at the N-terminus (Figure 3F). In addition, less abundant transcripts containing cryptic exon 3B that causes exon 4 of *BCL11B* to be read in a different ORF were detected with either viral or *BCL11B* encoded amino acids at the N-termini (Figures 3G and 3I).

To verify that the HPV70-*BCL11B* fusion RNAs were present, RT-PCR was performed on tumor RNA using one primer in virus sequences and the other in exon 4 (Figure 4A). This confirmed the presence of the fusion transcripts (Figure 4B). In addition to RNA from the tumor, a formalin fixed, paraffin embedded tissue fragment was obtained from a histopathology specimen from the same patient archived three years before tumor resection. The patient was diagnosed with CIN3 at that time, but was lost to follow-up. Both RNA and DNA were both prepared from the archived CIN3 sample. RT-PCR detected the presence of the fusion transcripts at both the tumor and the CIN3 stage (Figure 4B). Veracities of the RT-PCR products were confirmed by sequencing of excised bands from the tumor RT-PCR. The more prominent, faster migrating band confirmed splicing from HPV70 E1 to *BCL11B* exon 4 (**Supplementary Figure 4**). The faint upper band was also recovered from the tumor RT-PCR and sequenced thus confirming the structure of HPV70 E1 spliced to exon 3B, and exon 3B spliced to exon 4 of *BCL11B* in the same RNA (**Supplementary Figure 4**). Presence of the 3980 bp HPV70 DNA insertion into *BCL11B* was also confirmed by PCR testing for the HPV70 L1 junction in the CIN3 biopsy as well as in the tumor (Figure 4C). Thus, the HPV70 DNA insertion and fusion transcripts were present in the lesion by the time of CIN3 diagnosis as well as in the eventual tumor.

**Figure 4.**
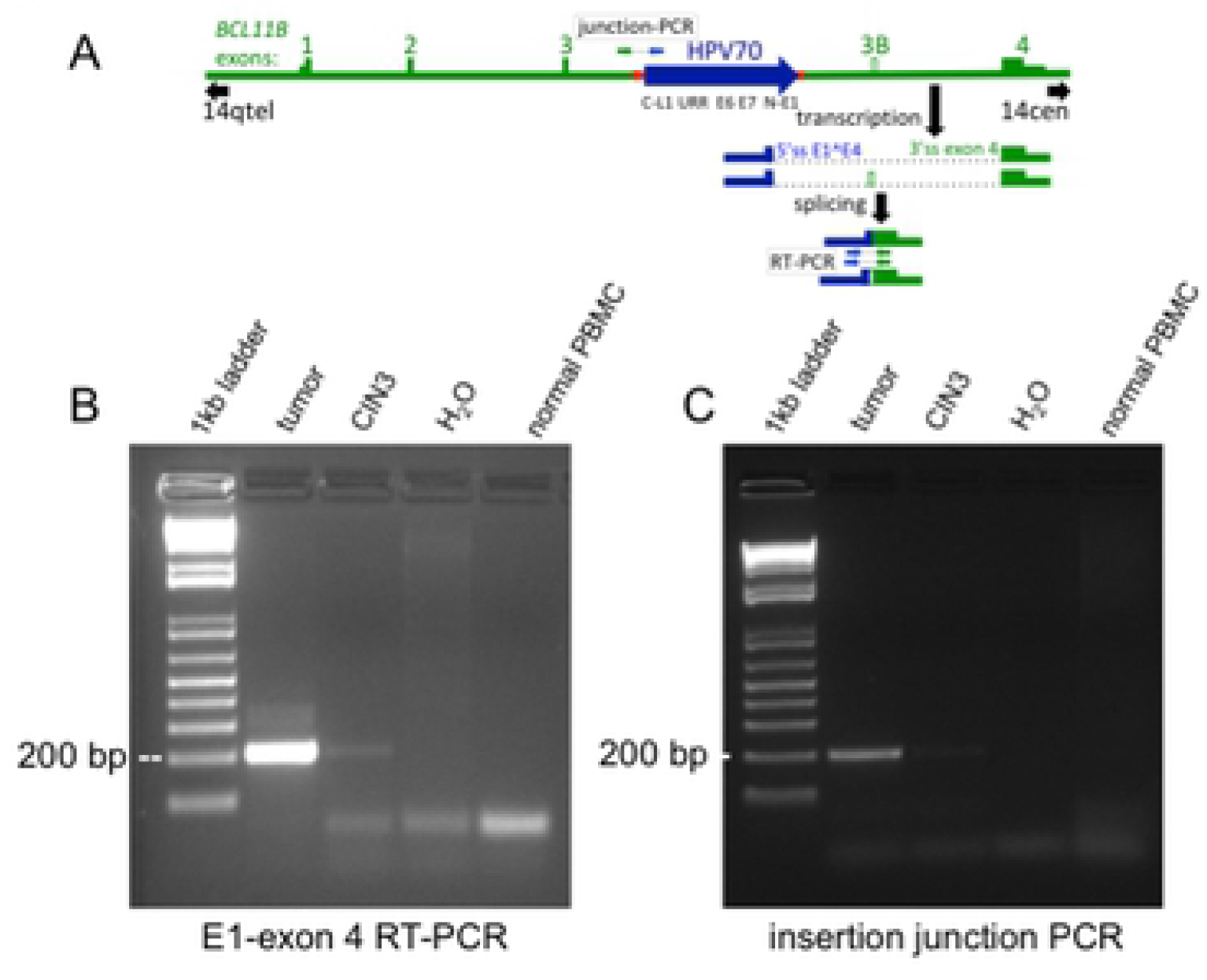
Detection of HPV70 – *BCL11B* RNA splicing and DNA insertion junctions in the tumor as well as in a CIN3 sample collected three years earlier from the same patient. **A)** Schematic diagram of the HPV70 DNA insert in *BCL11B* and the inferred transcripts with HPV70 in blue and *BCL11B* in green. Positions of the primers used for the junction PCR and the reverse transcriptase PCR (RT-PCR) are shown, along with the expected amplified products. **B)** RT-PCR confirming the HPV70 – exon 4 RNA splice junction in the tumor and detecting it in the CIN3 sample. Negative controls were H_2_O or PBMC RNA collected from a healthy donor. **C)** Junction PCR confirming the HPV70 DNA insertion in the tumor and detecting it in the CIN3 sample with comparable negative controls as in panel B. Sequences of the RT-PCR and junction PCR products are presented in Supplementary Figure 4.

## Discussion

Application of the HC+NGS approach to a cervical tumor identified the presence of HPV70 DNA and discerned the precise fraction of the viral genome present in the tumor. It also revealed that the viral DNA was integrated into one allele of *BCL11B* in the human genome, a gene that has been previously implicated in human tumorigenesis (43–47). Additional molecular genetic analyses identified the viral transcripts in the tumor and ascertained that the HPV70 DNA insertion resulted in fusion transcripts between the virus and portions of the *BCL11B* gene, including a novel cryptic exon. The finding of the viral insertion and the HPV70-*BCL11B* exon 4 fusion transcript in the CIN3 sample from the same patient indicated that these events had occurred by the intraepithelial precursor stage of disease and persisted during the invasive carcinoma stage. These results strongly imply that the HPV70 insertion and activation of *BCL11B* fusion ORF transcription were key events during malignant transformation. HPV70 was previously detected in ICCs and oral cancers (51, 52), and *BCL11B* was previously identified as a site of HPV16 integration in at least two cervical cancers (41, 42). Tumorigenesis by insertion of viral DNA near oncogenes is a thoroughly established oncogenic mechanism (31–35). The frequent detection of HPV DNA near oncogenes in virally induced human tumors indicates that insertional oncogenesis is a key mechanism in HPV-induced tumors, and that it may act in concert with HPV-encoded oncogenes, perhaps even replacing them in some instances (40). Detection of HPV70 DNA integrated into *BCL11B* and transcriptional activation of it, along with HPV E7 transcripts, strongly implicate HPV70 in this tumor.

Hybridization capture plus massively parallel sequencing has been used and proven to be reliable for identifying human genome integration sites of HPV DNA in tumors (20, 21, 24, 53). As demonstrated here, this approach is best applied by use of capture DNA probes encompassing the complete viral genomes of multiple HPV types. It allowed identification of the 3980 bp integrated HPV70 segment present in the tumor that comprised the URR-E6-E7-5’E1 region, indicating that this segment is the sole viral component responsible at least for maintaining the tumor state. Other viral components were required during at least the viral infection phase of tumorigenesis. HC+NGS also discerned the absence of the remainder of the viral genome in the tumor, including the 3’ portion of E1 and the entire E2 gene. Full-length viral E1 and E2 ORFs are frequently disrupted in ICCs (54). Precise mapping of the junctions at both ends of the integrated viral DNA, which can be achieved by HC+NGS, is required for full understanding of the consequences of an HPV DNA insertion. This includes the 34 bp human genome deletion observed here, and the presence of microhomologies at both ends of the inserted HPV70 segment suggesting that viral DNA integration presumably occurred by microhomology mediated repair (20). While the use of multiple analyses here revealed additional molecular insights, HC+NGS was sufficient to allow the identification of the HPV type in the tumor, inference of the viral DNA components present along with their organization, the exact location of the viral DNA in the human genome, and the inference of a specific human oncogene involved (*BCL11B*).

The WGS and SNP array analyses indicated that the HPV70 segment was integrated into one allele of *BCL11B*. Transcripts from the HPV70 segment encompassed the full-length E7 ORF of the viral E7 protein. This suggests that E7, BCL11B, and the chromosome 3q amplification were all likely important genetic components that function together at least to maintain this tumor. Surprisingly, the RNAseq results revealed that only a small fraction of transcripts encompassed the start codon of the E6 ORF, with most initiating further downstream in the E6 ORF. These findings suggest that only low levels of E6 might be translated in this tumor. In addition to binding to TP53, E6 also interacts with a number of anti-apoptotic proteins and PDZ domain-containing proteins (55–61). It also activates the *TERT* promoter (62–65). Perhaps *TERT* gene-containing chromosome 3q amplification in part compensated for the low levels of E6. The HPV70-*BCL11B* fusions that were detected in this tumor are strong candidates to function as dominant oncogenes in tumorigenesis. *BCL11B* encodes a Kruppel-like transcription factor with six C2H2-type zinc fingers. It has been implicated in both pediatric and adult T-cell acute lymphoblastic leukemia, where a high incidence of the t(5;14) chromosomal translocation and an association with expression of homeobox transcription factor TLX3 have been noted (43–47). BCL11B has been reported to be a component of the BAF (BRG1/BRM-associated factor) ATP-dependent chromatin remodeling complex. Genes encoding the various components of BAF are mutated in at least 20% of human cancers making it an attractive target for novel anti-tumor therapies such as inhibitors of the bromodomains that occur in other BAF components (66, 67). Most of the fusion ORFs identified here (Figure 3) encode an N-terminally truncated form of BCL11B that retains the six zinc finger domains. A small fraction encoded a form with a cryptic *BCL11B* exon that causes exon 4 to be read out of frame. It can be speculated that these altered forms of BCL11B perhaps have novel gain-of-function or dominant-negative function that alters transcription in a manner that drives tumorigenesis.

HPV tumorigenesis triggers genomic instability (21, 22). In addition to the chromosome 3q trisomy that recurrently occurs in ICC, copy number variation of multiple chromosome subregions was observed. Most striking was the *BCL11B* region of chromosome 14, with individual tumor cells displaying from two to over 50 copies. This observation stresses the perhaps surprising extent of genetic variability among individual tumor cells, and raises the question of what role it might play in tumorigenesis.

Population screening for HPV is based on demanding sensitivity and specificity criteria that balance clinical benefit with increased morbidity risk, emotional duress, and monetary costs of additional follow-up screening of individuals who test positive. While the limitations of screening tests for detection of HPV types that cause tumors of infrequent or unknown incidence have long been recognized, and broadly based HPV analyses of tumors have suggested roles for non-hrHPV’s in oncogenesis in a minority of tumors broader HPV type screening would create substantial burdens (1, 8, 68–70). Nonetheless, it is important to be aware that screening based solely on hrHPV detection may miss a small fraction of at risk individuals such as the patient described here, where the L1 gene PCR target may be deleted or may lack sufficient sequence homology for recognition. Expanded PCR specificity for HPV analysis is a subject of current interest (71).

HPV typing of ICCs currently offers little or no direct benefit to affected patients. The robust insights offered by the HC+NGS approach of simultaneous detection of a broad variety of HPV types, elucidation whether the viral DNA is integrated into the human genome, inference of the components of the viral genome that are present, and determination of whether the viral DNA is integrated adjacent to known human oncogenes suggest that it has the potential to provide deeper insights about individual tumors than simply assessing the presence of HPV DNA. One potential advantage of HC+NGS integration site analysis of human CIN lesions and tumors is determining whether HPV DNA integration in precancerous lesions has implications for progression to invasive carcinomas, including whether CIN lesions having HPV DNA integrations near known oncogenes warrant more intensive follow-up. A second is the identification of the full spectrum of human oncogenes where HPV DNAs are integrated in human tumors, perhaps in some instances providing actionable targets. A third is to provide a more complete global assessment of the frequencies of integrated DNA of HPV types currently considered to be of probable, possible, or low risk for tumorigenesis within tumors in various patient populations. Future studies could test if the mechanistic oncogenic insights gained from HC+NGS analyses have the ability to inform ongoing studies of disease management and perhaps lead to improved care.

## Methods

### Ethics Statement

Tumor and matching pretumor (CIN3) samples from a single patient were collected as part of an ongoing tissue collection protocol of the Department of Obstetrics & Gynecology and Women’s Health at the Albert Einstein College of Medicine. Tissue collection in adult patients (>18 years of age) occurred after written informed consent was obtained and subsequent experimental procedures were approved by the Internal Review Board (IRB) of the Albert Einstein College of Medicine (IRB#2009-265, IRB#2018-9256).

### Sample collection and nucleic acid preparation

As per tissue banking protocol, a fresh tumor biopsy sample was transported in sterile medium from the operating room to the Department of Pathology where sufficient tissue was procured for histopathologic diagnosis. The remaining tumor tissue was snap frozen in liquid nitrogen and transferred immediately to −80°C for storage until use. 10 µm sections of the frozen tissue were cut for H&E confirmation of the presence of tumor tissue by one of us who is a trained gynecologic pathologist (B.H.). DNA and RNA were extracted from sections of the frozen tissue using the QIAamp DNA Mini Kit and RNeasy Mini Kit (Qiagen), respectively, and stored at −80°C until use. CIN3 DNA and RNA were prepared using 10 µm sections from the patient’s archived CIN3 FFPE block and the RNeasy Plus Universal Kit (Qiagen), respectively. DNA and RNA concentrations were quantified using the Qubit Fluorometric Quantification method (Thermo Fisher Scientific) and their integrity assessed using the Bioanalyzer capillary electrophoresis system (Agilent).

### HPV DNA hybridization capture and short read Illumina sequencing

Targeted enrichment of HPV DNA was performed using hybridization capture probes homologous to the full-length genomes of 15 different HPV types (6, 11, 16, 18, 31, 33, 35, 39, 45, 52, 56, 58, 59, 68, 69) that were custom-designed using the Roche Nimblegen SeqCap EZ System (Roche, Basel, Switzerland). Biotinylated oligonucleotide probes specific for each HPV type were designed at ~150 bp intervals encompassing the positive strands of the complete ~8 kb double strand viral genomes to achieve 100% coverage. For library preparation, tumor genomic DNA was mechanically fragmented to 200 bp (Covaris, Woburn, MA), and Illumina adaptors were ligated at each end. Libraries were then hybridized to the custom HPV capture probes for 72 hours using the Roche target enrichment protocol following manufacture instructions and sequenced on one Illumina HiSeq 2500 lane (Illumina, San Diego, CA) using the paired end 150 bp sequencing mode. After sequencing, we aligned adaptor-cleaned, QC-passed, de-duplicated, paired end reads to a custom human (GRCh37/hg19) plus HPV reference genome containing 143 alpha genus HPV types from the Papillomavirus Episteme(72) using Burrows-Wheeler Aligner (BWA) (73). Junction fragments were computationally identified using a combination of programs; Delly (74) and SplazerS (75), both of which specify a combined read pair discordancy and split read analysis.

### Long sequencing reads using the Oxford Nanopore Technologies (ONT) MinION

Tumor-derived gDNA was purified and concentrated using the genomic DNA Clean and Concentrator-10 (gDCC-10) kit following manufacturer instructions (Zymo Research); concentration was assessed by Qubit and quality was assessed by Nanodrop (Thermo Fisher Scientific). The purified gDNA was then sheared by g-TUBE (Covaris, Woburn, MA) to 10 kb. Genomic libraries were prepared using the Oxford Nanopore 1D ligation library prep kit SQK-LSK108 following manufacturer instructions. Two independently prepared tumor DNA libraries were loaded onto two R9.4 flow cells and sequenced using a MinION Mk1b device (ONT) using the standard 48-hour scripts. Post-sequencing base calling and FASTQ extraction was performed using Albacore v2.0 (ONT), whereby raw voltage channel data is translated into canonical nucleotides. All libraries and quality filtered pass reads (Q≥7) were used for the subsequent analysis. Library-specific adaptors were trimmed and possible internal adaptors split using Porechop(76) with default parameters for adaptor identification (90% identity), end trimming (75% identity), and internal splitting (85% identity). Sequence reads were aligned to our custom combined reference genome using Ngmlr (77), a structural variant caller that uses a structural variant aware k-mer search to approximate alignments followed by a banded Smith-Waterman final alignment, along with a convex gap cost model to account for higher sequencing error frequencies associated with long reads. Structural variants were called using Sniffles (77) with parameter adjustment for expected low coverage.

### Detection of chromosome structural alterations, aneuploidy and copy number variation by whole genome sequencing and SNP analysis

For whole genome sequencing (WGS), 1 µg of genomic DNA from the tumor was used as input for library preparation using the NEBNext DNA Library Prep Kit (New England Biolabs). DNA was mechanically fragmented to 350 bp (Covaris, Woburn, MA) and the DNA fragments were end-polished, A-tailed, adaptor ligated, PCR amplified by P5 and indexed P7 oligos, and purified (AMPure XP system). The libraries were purified with AMPure XP (Beckman Coulter). QC passed libraries were sequenced on 2 Illumina HiSeq 2500 lanes to achieve 60X genome coverage on the paired end 150 bp mode. Reads with adaptor contamination or low quality (Phred Q30 <80%) were removed. After sequencing, paired end reads were aligned to our custom human (GRCh37/hg19) plus HPV70 reference genome using Burrows-Wheeler Aligner (BWA) (73). Files were mapped in BAM format, sorted using SAMtools(78), and duplicates removed using Picard (79). Genomic variant detection was accomplished for single nucleotide polymorphisms (SNPs)/small insertions and deletions (indels) using GATK v3.8 (80), structural variant (SV) detection using Delly v0.7.3 (74), and copy number variants (CNV) detection using control-FREEC (81). Following genomic variant detection, variants were annotated using the software ANNOVAR (82). Tumor genomic copy number was also assessed using the Genome-Wide Human SNP array 6.0 (Thermo Fisher Scientific). DNA prehandling and array hybridization were performed according to the manufacturer’s instructions (Affymetrix, Santa Clara, CA) and scanned in an Affymetrix GeneChip Scanner 3000. Data analysis and visualization was performed in Chromosomal Analysis Suite v4.0 (ChAS, Thermo Fisher Scientific) with a threshold of minimum 100 kb and 50 markers using NCBI build GRCh37/hg19.

### Global transcriptomic profiling of tumor RNA

Total RNA (2 µg) isolated from six, 10 µm, tumor sections with an RNA integrity number of 9.3 (Agilent 1000) and subjected to oligoT magnetic bead enrichment. Libraries were prepared following the NEB standard protocol as follows: cDNA was synthesized using random hexamer primers and M-MuLV RT (RNaseH-) followed by second strand synthesis with DNA polymerase I and RNaseH. The double stranded cDNA was purified using AMPure XP beads (Beckman Coulter), end tail repaired, adaptor ligated, size-selected, and PCR amplified before Illumina sequencing. 150 bp insert cDNA libraries were sequenced to a depth of 60 million reads on one Illumina HiSeq 2500 lane (Illumina, Inc., San Diego, CA) using the paired end 150 bp mode. Raw image data were transformed to sequenced reads by CASAVA and stored in FASTQ format. Raw reads were then filtered to remove adaptors, reads containing N>10% (N representing undetermined bases), and reads of Qscore (Quality value) of >50% of bases ≤ 5. Of approximately 144 million reads, 97.8% passed filtering with >96% having Phred scores >30. Gene model annotation files were downloaded from the genome website browser (NCBI/UCSC/Ensembl) for (GRCh37/hg19) and in custom format from Papillomavirus Episteme (72) for HPV70. Indexes of the custom reference genome were built, and paired-end clean reads were aligned to the reference using STAR v2.5 (83). HTSeq v0.6.1 (84) was used to count the read numbers mapped of each gene. As there was no tumor-free matched control for this patient, differential expression analysis was not performed. As our main interest was in potential fusion transcripts between *BCL11B* and HPV70, STAR-Fusion 0.8.0 (85) was used for the detection of fusion transcripts.

### Validation of HPV DNA integration junctions & HPV70-*BCL11B* Fusion Transcripts

To detect the integrated HPV70 DNA segment, PCR primers were designed to flank each side of the HPV70 genome – human genome junctions obtained from HC+NGS split reads. The chr14:99,683,287 – HPV70 L1:6306 primers were 5’-TTCCAAAAGTGTCTGGCAAA and 5’GTGTGTGAATGTGGGGGTGT producing a 216 bp amplicon. The chr14:99683253 – HPV70 E1:2380 primer pair was 5’-TACAGGGACCACCAAACACA and 5’-CTGGACTGCACACAGACACA producing a 168 bp amplicon, and PCR reactions were performed using GoTaq Green (Promega) using 30 cycles of 94°C for 30 sec, 60°C for 30 sec, and 72°C for 45 sec. To detect the HPV70-*BCL11B* fusion transcripts, primers were designed to flank each side of the fusion splice junction from the HPV70 E1 5’ss at genome position 943 to the *BCL11B* exon 4 3’ss at GRCh37/hg19 genome position chr14:99642532 obtained from RNAseq split reads. The HPV70 primer was 5’-GAAGAACCACAGCGTCACAA and the *BCL11B* exon 4 primer was 5’-TGCAAATGTAGCTGGAAGGC with a 214 bp amplicon. First-strand cDNA synthesis was performed on 5 µg of total RNA from either fresh tumor tissue or archival FFPE CIN3 tissue using the exon 4 primer and SuperScript™II RT (Thermo Fisher Scientific) following manufacture’s protocol. The subsequent PCR to detect the presence of the fusion transcripts was performed using 35 cycles of 95°C for 15 sec, 60°C for 30 sec, and 72°C for 45 sec. The resultant PCR products were resolved on 0.8% or 2.0% agarose gels, respectively, and the bands were excised using the Monarch DNA Gel Extraction Kit (New England Biolabs) for Sanger sequencing analysis.

### *In situ* visualization of *BCL11B* locus

Fluorescence in situ hybridization (FISH) was used to visualize the *BCL11B* at the single cell level on histopathologic sections of the tumor. BAC RP-11-179F4 spanning the *BCL11B* locus was obtained from the BACPAC Resources Center at the Children’s Hospital Oakland Research Institute. 8 µm sections were cut from the diagnostic FFPE tumor block from the patient, and mounted on positively charged slides. Slides were incubated overnight at 56°C prior to FISH hybridization that was performed as previously described (86). Briefly, the slides were deparaffinized in Hemo-De at room temperature for 10 minutes × 2, dehydrated in 100% ethanol for 5 minutes × 2 and placed on a 50°C slide warmer for 5 minutes. The slides were then pretreated for 24 minutes using the Vysis Paraffin Pretreatment Reagent Kit (Abbott Molecular), and fixed in 10% buffered formalin as per the protocol. The FISH probe was labeled by nick translation using DY-415-aadUTP (Dyomics, Jena, GE, USA) as previously described (86) and hybridized overnight. The slides were then washed in 0.4X SSC pre-warmed to 74°C, followed by 4X SSC/0.1% Tween. Images were acquired with a manual inverted fluorescence microscope (Axiovert 200, Zeiss) with fine focusing oil immersion lens (x40, NA 1.3). The resulting emissions were collected using 350-to-470 nm (for DAPI) and 470-to-540 nm (for DY-495-dUTP) filters. The microscope was equipped with a Camera Hall 100 and the Applied Spectral Imaging software. Images representing 90 nuclei were randomly acquired and saved as. tiff composite files. Single cell copy numbers of the *BCL11B* locus were manually counted and categorized as described in the figures.

## Acknowledgments

We thank Rob Coleman, Matt Gamble, Charles Query, and Deyou Zhang for helpful discussions. We thank the Molecular Cytogenetic Core at Albert Einstein College of Medicine, and in particular Jidong Shan and Yinghui Song for assisting with the FISH studies. This research was supported by the Albert Einstein Cancer Center Support Grant of the National Institutes of Health under award number P30CA013330 (C.M. and J.L.), by The American Association of Obstetricians and Gynecologists Foundation and The American Board of Obstetrics and Gynecology scholar award (A.V.), and The Foundation for Women’s Cancer 2016 ME STRONG Young Investigator’s Award (A.V.).

## Author Contributions

AVA performed and supervised the experiments NEP, ECM performed the experiments

LA, KVD contributed intellectually to the experimental design BH performed the histopathological analysis

NN, DYK, MH contributed intellectually to the project design AVA, JL and CM interpreted the results

JL and CM supervised the entire project

All authors contributed to writing the manuscript

**Supplementary Figure 1.** Sequence of the HPV70 reference genome is presented. *‘s indicate the nucleotide was present in the viral DNA insertion in the tumor, and that the nucleotide was identical. Positions where the tumor nucleotides differed are highlighted in yellow and the change is shown. The gray highlighted segment is the URR between the L1 and E6 ORFs.

**Supplementary Figure 2.** Five of the split reads crossing the HPV70 E1^E4 5’ss to *BCL11B* exon 4 splice junction. HPV70 sequences are highlighted in blue. Human genome *BCL11B* sequences are highlighted in green.

**Supplementary Figure 3. (3A)** DNA sequence of *BCL11B* exon 3B (upper case letters, hg19 position chr14:99665867 to 99665710) with immediately flanking intron sequences (lower case letters). Upstream intron sequences include 3’ss consensus sequence (bold), t-rich segment, and potential lariat site (ac). Downstream intron sequences include 5’ss consensus (bold). (**3B**) The four RNAseq split reads crossing the HPV70 E1^E4 5’ss to *BCL11B* exon 3B splice junction. HPV70 sequences are highlighted in blue. Human genome *BCL11B* sequences are highlighted in green.

**Supplementary Figure 4. (4A)** Chromatograms and sequences of RT-PCR products of the RT-PCR shown in Figure 4B highlighting the HPV70 (blue), *BCL11B* (green), and micro-homology overlap (red) sequences. **(4B)** Chromatograms and sequences of RT-PCR products shown in Figure 4C highlighting the HPV70 (blue), *BCL11B* exon 3B (gray), and *BCL11B* exon 4 (green) sequences of the PCR products generated from tumor RNA.

## References

1. Walboomers JM, Jacobs MV, Manos MM, Bosch FX, Kummer JA, Shah KV, et al. Human papillomavirus is a necessary cause of invasive cervical cancer worldwide. J Pathol. 1999;189(1):12–9.

2. Gaglia MM, Munger K. More than just oncogenes: mechanisms of tumorigenesis by human viruses. Curr Opin Virol. 2018;32:48–59.

3. Li N, Franceschi S, Howell-Jones R, Snijders PJ, Clifford GM. Human papillomavirus type distribution in 30,848 invasive cervical cancers worldwide: Variation by geographical region, histological type and year of publication. Int J Cancer. 2011;128(4):927–35.

4. Hopenhayn C, Christian A, Christian WJ, Watson M, Unger ER, Lynch CF, et al. Prevalence of human papillomavirus types in invasive cervical cancers from 7 US cancer registries before vaccine introduction. J Low Genit Tract Dis. 2014;18(2):182–9.

5. Bouvard V, Baan R, Straif K, Grosse Y, Secretan B, El Ghissassi F, et al. A review of human carcinogens--Part B: biological agents. Lancet Oncol. 2009;10(4):321–2.

6. Prigge ES, von Knebel Doeberitz M, Reuschenbach M. Clinical relevance and implications of HPV-induced neoplasia in different anatomical locations. Mutat Res Rev Mutat Res. 2017;772:51–66.

7. Munoz N, Bosch FX, de Sanjose S, Herrero R, Castellsague X, Shah KV, et al. Epidemiologic classification of human papillomavirus types associated with cervical cancer. N Engl J Med. 2003;348(6):518–27.

8. Walboomers JM, Meijer CJ. Do HPV-negative cervical carcinomas exist? J Pathol. 1997;181(3):253–4.

9. Ronco G, Dillner J, Elfstrom KM, Tunesi S, Snijders PJ, Arbyn M, et al. Efficacy of HPV-based screening for prevention of invasive cervical cancer: follow-up of four European randomised controlled trials. Lancet. 2014;383(9916):524–32.

10. Rijkaart DC, Berkhof J, van Kemenade FJ, Coupe VM, Rozendaal L, Heideman DA, et al. HPV DNA testing in population-based cervical screening (VUSA-Screen study): results and implications. Br J Cancer. 2012;106(5):975–81.

11. Leinonen MK, Nieminen P, Lonnberg S, Malila N, Hakama M, Pokhrel A, et al. Detection rates of precancerous and cancerous cervical lesions within one screening round of primary human papillomavirus DNA testing: prospective randomised trial in Finland. BMJ. 2012;345:e7789.

12. Wright TC, Stoler MH, Behrens CM, Sharma A, Zhang G, Wright TL. Primary cervical cancer screening with human papillomavirus: end of study results from the ATHENA study using HPV as the first-line screening test. Gynecol Oncol. 2015;136(2):189–97.

13. Ogilvie GS, van Niekerk D, Krajden M, Smith LW, Cook D, Gondara L, et al. Effect of Screening With Primary Cervical HPV Testing vs Cytology Testing on High-grade Cervical Intraepithelial Neoplasia at 48 Months: The HPV FOCAL Randomized Clinical Trial. JAMA. 2018;320(1):43–52.

14. Schiffman M, Khan MJ, Solomon D, Herrero R, Wacholder S, Hildesheim A, et al. A study of the impact of adding HPV types to cervical cancer screening and triage tests. J Natl Cancer Inst. 2005;97(2):147–50.

15. Koutsky L. Epidemiology of genital human papillomavirus infection. Am J Med. 1997;102(5A):3–8.

16. Corden SA, Sant-Cassia LJ, Easton AJ, Morris AG. The integration of HPV-18 DNA in cervical carcinoma. Mol Pathol. 1999;52(5):275–82.

17. Klaes R, Woerner SM, Ridder R, Wentzensen N, Duerst M, Schneider A, et al. Detection of high-risk cervical intraepithelial neoplasia and cervical cancer by amplification of transcripts derived from integrated papillomavirus oncogenes. Cancer Res. 1999;59(24):6132–6.

18. Schwarz E, Freese UK, Gissmann L, Mayer W, Roggenbuck B, Stremlau A, et al. Structure and transcription of human papillomavirus sequences in cervical carcinoma cells. Nature. 1985;314(6006):111–4.

19. Einstein MH, Cruz Y, El-Awady MK, Popescu NC, DiPaolo JA, van Ranst M, et al. Utilization of the human genome sequence localizes human papillomavirus type 16 DNA integrated into the TNFAIP2 gene in a fatal cervical cancer from a 39-year-old woman. Clin Cancer Res. 2002;8(2):549–54.

20. Hu Z, Zhu D, Wang W, Li W, Jia W, Zeng X, et al. Genome-wide profiling of HPV integration in cervical cancer identifies clustered genomic hot spots and a potential microhomology-mediated integration mechanism. Nat Genet. 2015;47(2):158–63.

21. Holmes A, Lameiras S, Jeannot E, Marie Y, Castera L, Sastre-Garau X, et al. Mechanistic signatures of HPV insertions in cervical carcinomas. NPJ Genom Med. 2016;1:16004.

22. Akagi K, Li J, Broutian TR, Padilla-Nash H, Xiao W, Jiang B, et al. Genome-wide analysis of HPV integration in human cancers reveals recurrent, focal genomic instability. Genome Res. 2014;24(2):185–99.

23. Xu B, Chotewutmontri S, Wolf S, Klos U, Schmitz M, Durst M, et al. Multiplex Identification of Human Papillomavirus 16 DNA Integration Sites in Cervical Carcinomas. PLoS One. 2013;8(6):e66693.

24. Liu Y, Lu Z, Xu R, Ke Y. Comprehensive mapping of the human papillomavirus (HPV) DNA integration sites in cervical carcinomas by HPV capture technology. Oncotarget. 2016;7(5):5852–64.

25. Zhang R, Shen C, Zhao L, Wang J, McCrae M, Chen X, et al. Dysregulation of host cellular genes targeted by human papillomavirus (HPV) integration contributes to HPV-related cervical carcinogenesis. Int J Cancer. 2016;138(5):1163–74.

26. Kumar Gupta A, Kumar M. HPVbase--a knowledgebase of viral integrations, methylation patterns and microRNAs aberrant expression: As potential biomarkers for Human papillomaviruses mediated carcinomas. Sci Rep. 2015;5:12522.

27. Schaeffer AJ, Nguyen M, Liem A, Lee D, Montagna C, Lambert PF, et al. E6 and E7 oncoproteins induce distinct patterns of chromosomal aneuploidy in skin tumors from transgenic mice. Cancer Res. 2004;64(2):538–46.

28. Kadaja M, Isok-Paas H, Laos T, Ustav E, Ustav M. Mechanism of genomic instability in cells infected with the high-risk human papillomaviruses. PLoS Pathog. 2009;5(4):e1000397.

29. Peter M, Stransky N, Couturier J, Hupe P, Barillot E, de Cremoux P, et al. Frequent genomic structural alterations at HPV insertion sites in cervical carcinoma. J Pathol. 2010;221(3):320–30.

30. Jang MK, Shen K, McBride AA. Papillomavirus genomes associate with BRD4 to replicate at fragile sites in the host genome. PLoS Pathog. 2014;10(5):e1004117.

31. Neel BG, Hayward WS, Robinson HL, Fang J, Astrin SM. Avian leukosis virus-induced tumors have common proviral integration sites and synthesize discrete new RNAs: oncogenesis by promoter insertion. Cell. 1981;23(2):323–34.

32. Payne GS, Bishop JM, Varmus HE. Multiple arrangements of viral DNA and an activated host oncogene in bursal lymphomas. Nature. 1982;295(5846):209–14.

33. Suzuki T, Shen H, Akagi K, Morse HC, Malley JD, Naiman DQ, et al. New genes involved in cancer identified by retroviral tagging. Nat Genet. 2002;32(1):166–74.

34. Mikkers H, Allen J, Knipscheer P, Romeijn L, Hart A, Vink E, et al. High-throughput retroviral tagging to identify components of specific signaling pathways in cancer. Nat Genet. 2002;32(1):153–9.

35. Lund AH, Turner G, Trubetskoy A, Verhoeven E, Wientjens E, Hulsman D, et al. Genome-wide retroviral insertional tagging of genes involved in cancer in Cdkn2a-deficient mice. Nat Genet. 2002;32(1):160–5.

36. Hacein-Bey-Abina S, Garrigue A, Wang GP, Soulier J, Lim A, Morillon E, et al. Insertional oncogenesis in 4 patients after retrovirus-mediated gene therapy of SCID-X1. J Clin Invest. 2008;118(9):3132–42.

37. Braun CJ, Boztug K, Paruzynski A, Witzel M, Schwarzer A, Rothe M, et al. Gene therapy for Wiskott-Aldrich syndrome--long-term efficacy and genotoxicity. Sci Transl Med. 2014;6(227):227ra33.

38. Warburton A, Redmond CJ, Dooley KE, Fu H, Gillison ML, Akagi K, et al. HPV integration hijacks and multimerizes a cellular enhancer to generate a viral-cellular super-enhancer that drives high viral oncogene expression. PLoS Genet. 2018;14(1):e1007179.

39. McBride AA, Warburton A. The role of integration in oncogenic progression of HPV-associated cancers. PLoS Pathog. 2017;13(4):e1006211.

40. Yuan H, Krawczyk E, Blancato J, Albanese C, Zhou D, Wang N, et al. HPV positive neuroendocrine cervical cancer cells are dependent on Myc but not E6/E7 viral oncogenes. Sci Rep. 2017;7:45617.

41. Ojesina AI, Lichtenstein L, Freeman SS, Pedamallu CS, Imaz-Rosshandler I, Pugh TJ, et al. Landscape of genomic alterations in cervical carcinomas. Nature. 2014;506(7488):371–5.

42. Li H, Yang Y, Zhang R, Cai Y, Yang X, Wang Z, et al. Preferential sites for the integration and disruption of human papillomavirus 16 in cervical lesions. J Clin Virol. 2013;56(4):342–7.

43. Berger R, Dastugue N, Busson M, Van Den Akker J, Perot C, Ballerini P, et al. t(5;14)/HOX11L2-positive T-cell acute lymphoblastic leukemia. A collaborative study of the Groupe Francais de Cytogenetique Hematologique (GFCH). Leukemia. 2003;17(9):1851–7.

44. Cave H, Suciu S, Preudhomme C, Poppe B, Robert A, Uyttebroeck A, et al. Clinical significance of HOX11L2 expression linked to t(5;14)(q35;q32), of HOX11 expression, and of SIL-TAL fusion in childhood T-cell malignancies: results of EORTC studies 58881 and 58951. Blood. 2004;103(2):442–50.

45. MacLeod RA, Nagel S, Drexler HG. BCL11B rearrangements probably target T-cell neoplasia rather than acute myelocytic leukemia. Cancer Genet Cytogenet. 2004;153(1):88–9.

46. Wakabayashi Y, Inoue J, Takahashi Y, Matsuki A, Kosugi-Okano H, Shinbo T, et al. Homozygous deletions and point mutations of the Rit1/Bcl11b gene in gamma-ray induced mouse thymic lymphomas. Biochem Biophys Res Commun. 2003;301(2):598–603.

47. Oshiro A, Tagawa H, Ohshima K, Karube K, Uike N, Tashiro Y, et al. Identification of subtype-specific genomic alterations in aggressive adult T-cell leukemia/lymphoma. Blood. 2006;107(11):4500–7.

48. Heselmeyer K, Schrock E, du Manoir S, Blegen H, Shah K, Steinbeck R, et al. Gain of chromosome 3q defines the transition from severe dysplasia to invasive carcinoma of the uterine cervix. Proc Natl Acad Sci U S A. 1996;93(1):479–84.

49. Heselmeyer-Haddad K, Sommerfeld K, White NM, Chaudhri N, Morrison LE, Palanisamy N, et al. Genomic amplification of the human telomerase gene (TERC) in pap smears predicts the development of cervical cancer. Am J Pathol. 2005;166(4):1229–38.

50. Stacey SN, Jordan D, Williamson AJ, Brown M, Coote JH, Arrand JR. Leaky scanning is the predominant mechanism for translation of human papillomavirus type 16 E7 oncoprotein from E6/E7 bicistronic mRNA. J Virol. 2000;74(16):7284–97.

51. Nielsen A, Iftner T, Norgaard M, Munk C, Junge J, Kjaer SK. The importance of low-risk HPV infection for the risk of abnormal cervical cytology/histology in more than 40 000 Danish women. Sex Transm Infect. 2012;88(8):627–32.

52. Pannone G, Santoro A, Carinci F, Bufo P, Papagerakis SM, Rubini C, et al. Double demonstration of oncogenic high risk human papilloma virus DNA and HPV-E7 protein in oral cancers. Int J Immunopathol Pharmacol. 2011;24(2 Suppl):95–101.

53. Liu Y, Zhang C, Gao W, Wang L, Pan Y, Gao Y, et al. Genome-wide profiling of the human papillomavirus DNA integration in cervical intraepithelial neoplasia and normal cervical epithelium by HPV capture technology. Sci Rep. 2016;6:35427.

54. Cancer Genome Atlas Research N, Albert Einstein College of M, Analytical Biological S, Barretos Cancer H, Baylor College of M, Beckman Research Institute of City of H, et al. Integrated genomic and molecular characterization of cervical cancer. Nature. 2017;543(7645):378–84.

55. Wallace NA, Galloway DA. Novel Functions of the Human Papillomavirus E6 Oncoproteins. Annu Rev Virol. 2015;2(1):403–23.

56. Galloway DA, Laimins LA. Human papillomaviruses: shared and distinct pathways for pathogenesis. Curr Opin Virol. 2015;14:87–92.

57. Harden ME, Munger K. Perturbation of DROSHA and DICER expression by human papillomavirus 16 oncoproteins. Virology. 2017;507:192–8.

58. Chiang C, Pauli EK, Biryukov J, Feister KF, Meng M, White EA, et al. The Human Papillomavirus E6 Oncoprotein Targets USP15 and TRIM25 To Suppress RIG-I-Mediated Innate Immune Signaling. J Virol. 2018;92(6).

59. Westrich JA, Warren CJ, Pyeon D. Evasion of host immune defenses by human papillomavirus. Virus Res. 2017;231:21–33.

60. Gunasekharan VK, Li Y, Andrade J, Laimins LA. Post-Transcriptional Regulation of KLF4 by High-Risk Human Papillomaviruses Is Necessary for the Differentiation-Dependent Viral Life Cycle. PLoS Pathog. 2016;12(7):e1005747.

61. Wallace NA, Khanal S, Robinson KL, Wendel SO, Messer JJ, Galloway DA. High-Risk Alphapapillomavirus Oncogenes Impair the Homologous Recombination Pathway. J Virol. 2017;91(20).

62. Zhang Y, Dakic A, Chen R, Dai Y, Schlegel R, Liu X. Direct HPV E6/Myc interactions induce histone modifications, Pol II phosphorylation, and hTERT promoter activation. Oncotarget. 2017;8(56):96323–39.

63. Klingelhutz AJ, Foster SA, McDougall JK. Telomerase activation by the E6 gene product of human papillomavirus type 16. Nature. 1996;380(6569):79–82.

64. Veldman T, Horikawa I, Barrett JC, Schlegel R. Transcriptional activation of the telomerase hTERT gene by human papillomavirus type 16 E6 oncoprotein. J Virol. 2001;75(9):4467–72.

65. Gewin L, Galloway DA. E box-dependent activation of telomerase by human papillomavirus type 16 E6 does not require induction of c-myc. J Virol. 2001;75(15):7198–201.

66. Kadoch C, Crabtree GR. Mammalian SWI/SNF chromatin remodeling complexes and cancer: Mechanistic insights gained from human genomics. Sci Adv. 2015;1(5):e1500447.

67. Hodges C, Kirkland JG, Crabtree GR. The Many Roles of BAF (mSWI/SNF) and PBAF Complexes in Cancer. Cold Spring Harb Perspect Med. 2016;6(8).

68. Halec G, Alemany L, Lloveras B, Schmitt M, Alejo M, Bosch FX, et al. Pathogenic role of the eight probably/possibly carcinogenic HPV types 26, 53, 66, 67, 68, 70, 73 and 82 in cervical cancer. J Pathol. 2014;234(4):441–51.

69. Arbyn M, Tommasino M, Depuydt C, Dillner J. Are 20 human papillomavirus types causing cervical cancer? J Pathol. 2014;234(4):431–5.

70. Halec G, Alemany L, Quiros B, Clavero O, Hofler D, Alejo M, et al. Biological relevance of human papillomaviruses in vulvar cancer. Mod Pathol. 2017;30(4):549–62.

71. Demarco M, Carter-Pokras O, Hyun N, Castle PE, He X, Dallal CM, et al. Validation of a Human Papillomavirus (HPV) DNA Cervical Screening Test That Provides Expanded HPV Typing. J Clin Microbiol. 2018;56(5).

72. Papillomavirus Episteme [Available from: http://pave.niaid.nih.gov.

73. Li H, Durbin R. Fast and accurate short read alignment with Burrows-Wheeler transform. Bioinformatics. 2009;25(14):1754–60.

74. Rausch T, Zichner T, Schlattl A, Stutz AM, Benes V, Korbel JO. DELLY: structural variant discovery by integrated paired-end and split-read analysis. Bioinformatics. 2012;28(18):i333–i9.

75. Emde AK, Schulz MH, Weese D, Sun R, Vingron M, Kalscheuer VM, et al. Detecting genomic indel variants with exact breakpoints in single- and paired-end sequencing data using SplazerS. Bioinformatics. 2012;28(5):619–27.

76. Porechop [Available from: https://github.com/rrwick/Porechop.

77. Sedlazeck FJ, Rescheneder P, Smolka M, Fang H, Nattestad M, von Haeseler A, et al. Accurate detection of complex structural variations using single-molecule sequencing. Nat Methods. 2018;15(6):461–8.

78. Li H, Handsaker B, Wysoker A, Fennell T, Ruan J, Homer N, et al. The Sequence Alignment/Map format and SAMtools. Bioinformatics. 2009;25(16):2078–9.

79. Picard.

80. DePristo MA, Banks E, Poplin R, Garimella KV, Maguire JR, Hartl C, et al. A framework for variation discovery and genotyping using next-generation DNA sequencing data. Nat Genet. 2011;43(5):491–8.

81. Boeva V, Popova T, Bleakley K, Chiche P, Cappo J, Schleiermacher G, et al. Control-FREEC: a tool for assessing copy number and allelic content using next-generation sequencing data. Bioinformatics. 2012;28(3):423–5.

82. Wang K, Li M, Hakonarson H. ANNOVAR: functional annotation of genetic variants from high-throughput sequencing data. Nucleic Acids Res. 2010;38(16):e164.

83. Dobin A, Davis CA, Schlesinger F, Drenkow J, Zaleski C, Jha S, et al. STAR: ultrafast universal RNA-seq aligner. Bioinformatics. 2013;29(1):15–21.

84. Anders S, Pyl PT, Huber W. HTSeq--a Python framework to work with high-throughput sequencing data. Bioinformatics. 2015;31(2):166–9.

85. Haas BJ DA, Stransky N, Li B, Yang X, Tickle T, Bankapur A, Ganote C, Doak TG, Pochet N, Sun J, Wu CJ, Gingeras TR, Regev A. STAR-Fusion: Fast and Accurate Fusion Transcript Detection from RNA-seq. bioRxiv [Internet]. 120295. Available from: https://doi.org/10.1101/120295.

86. Andriani GA, Almeida VP, Faggioli F, Mauro M, Tsai WL, Santambrogio L, et al. Whole Chromosome Instability induces senescence and promotes SASP. Sci Rep. 2016;6:35218.

